# Vocal-visual combinations in wild chimpanzees

**DOI:** 10.1101/2023.05.24.541909

**Authors:** Joseph G. Mine, Claudia Wilke, Chiara Zulberti, Melika Bejhati, Sabine Stoll, Zarin Machanda, Andri Manser, Katie. E Slocombe, Simon W. Townsend

## Abstract

Human communication is strikingly multi-modal, relying on vocal utterances combined with visual gestures, facial expressions and more. Recent efforts to describe multi-modal signal production in our ape relatives have shed important light on the evolutionary trajectory of this core hallmark of human language. However, whilst promising, a systematic quantification of primate signal production which filters out random combinations produced across modalities is currently lacking. Here, through recording the communicative behaviour of wild chimpanzees from the Kibale forest, Uganda we address this issue and generate the first repertoire of non-random combined vocal and visual components. Using collocation analysis, we identify more than 100 vocal-visual combinations which occur more frequently than expected by chance. We also probe how multi-modal production varies in the population, finding no differences between individuals as a function of age, sex or rank. The number of visual components exhibited alongside vocalizations was, however, associated with vocalization type and duration. We demonstrate that chimpanzees produce a vast array of combined vocal and visual components, exhibiting a hitherto underappreciated level of combinatorial complexity. We conclude that a multi-modal approach is crucial to accurately representing the communicative abilities of non-human primates.

## Introduction

Human communication, which is crucial to our daily lives, is an inherently multi-component system [1]. When speaking, humans typically accompany their utterances with gestures, facial expressions and other signals or cues. A smile, for example, or a shrug, may enhance the meaning of an utterance and influence the receiver’s interpretation [2]. The combination of vocal utterances with such additional cues, known as extralinguistic cues (ELCs) [3], allows speakers to convey rich and multifaceted meanings and is therefore arguably a cornerstone of the human language faculty [4]. Whether similar multi-modal signals are employed in the communication systems of non-human primates has received growing attention, given the valuable insight such data can provide regarding the evolutionary origins of human communication and language [5,6]. The term “multi-modal” has, however, been used differently in previous communication studies, in some cases denoting multiple signaling channels (e.g. facial expressions vs gestures) [7,8], while in others denoting multiple sensory modalities (e.g. acoustic vs visual modality) [9,10]. Here, we define a multi-modal signal as one that is received in at least two sensory modalities. Previous research in non-human primate communication has shown that apes augment their vocalizations with specific visual gestures, potentially as a way to disambiguate or refine meaning, akin to the function of extralinguistic cues as semantic devices in language [8,11]. For example, in bonobos, the “contest hoot” vocalization can be combined with a threatening “stomp” gesture during agonistic challenges, or with a playful “wrist shake” in friendly play [12]. Similarly, in chimpanzees, mothers interacting with infants often combine the “soft hoo” vocalization with the “arm reach” or “present back” gesture, to invite the infant to climb onto their back [13].

To date, the most thorough attempt to document multi-modal signal production in apes has established a repertoire of combinations of existing vocalizations, gestures and facial expressions in chimpanzees [8]. However, since vocalizations may co-occur with other signals or cues simply by chance, differentiating random from non-random multi-modal combinations is a critical step, ultimately providing a more accurate reflection of the multi-modal proclivities of a species. Such a data-driven quantification of the vocal-visual repertoire is currently lacking for any primate [5,6]. We aimed to bridge this gap in understanding through systematically investigating the multi-modal communicative behaviour of wild chimpanzees. As a first step, we build a vocal-visual repertoire by focusing on naturally occurring vocal production and recording the accompanying visual components. Through applying methods borrowed from computational linguistics, namely collocation analysis, we then quantify the non-random nature of identified vocal-visual combinations [14].

Chimpanzees, like humans, have complex social lives: they reside in groups of ∼50-100 individuals, forming strong and durable relationships with relatives as well as non-kin [15]. Likely as a way to navigate this complex social environment, chimpanzees are also equipped with a rich system of communication comprising signals and cues from both visual and vocal modalities [16-18]. The vocal repertoire consists of approximately 13 different call types [16]. The repertoire is commonly described as graded, meaning that there is acoustic variation within a single category, as well as a degree of overlap in acoustic features also between certain categories. The anatomy of the chimpanzee brain and vocal tract constrains vocal production to a limited range of sounds compared to human vocal production [19,20]. By contrast, visual signal production in chimpanzees is highly flexible and the repertoire is vast, comprising at least 9 facial expressions [18] and 66 gesture types [17].

Importantly, vocal signals, facial expressions and manual gestures are complemented by an equally broad array of body movements or behaviours, which might be rather described as cues (i.e. behaviours that have not necessarily evolved for a communicative purpose, yet may carry some communicative value) [21,22]. For example, a chimpanzee’s body posture (e.g. sitting vs standing), or the orientation of their gaze, which can be towards or away from the recipient, may carry important communicative value for the recipient. As such, we adopted an inclusive, bottom-up approach and considered the combination of vocal signals with both visual signals and behaviours that may act as cues. To this end, we recorded all visible movements, body postures, orientations, behaviours, gestures or facial expressions exhibited by the signaler alongside the vocalization as non-vocal behaviours (NVBs).

In addition to establishing a repertoire of non-random vocal-visual combinations, we aimed to examine the variation underlying NVB production within the population. Previous research has implicated various demographic factors, such as age, sex and rank in driving variation in both gestural and vocal behaviour. For example, females are known to produce a higher rate of call combinations than their male counterparts [23], while highest-ranking males were shown to be the most prolific gesture producers [11]. In line with this existing body of work we therefore also probed how demographic factors influenced the combination of visual components with vocal signals. Given our data-driven and exploratory approach, we formulate no *a priori* predictions regarding patterns of demographic variation. Finally, we probe how NVB production changes in accordance with the characteristics of the call. For example, calls produced while feeding may be associated with different amounts of NVBs compared to calls produced upon encounters with conspecifics. In addition, call duration might affect NVB production as longer calls might be associated with more movements, changes in body posture or gestures. Therefore, we test whether NVB production is influenced by call type and duration.

## Methods

### Study site and data collection

The study was conducted on wild chimpanzees from the Kanyawara community in Kibale national park, Uganda [24]. The population consists of ∼60 individuals inhabiting a home range of ∼15km^2^. The Kanyawara community has been the object of long-term study since 1987 and is entirely habituated. The data used in this study were collected between February-May 2013, and between June 2014 and March 2015 [8]. These data consist in video-audio recordings collected within the chimpanzee home range, between 0800 and 1900 hours. The equipment included a hand-held camcorder (Panasonic HDC-SD90), and an external microphone (Sennheiser MKE 400).

The individuals observed in this study were 13 females and 14 males, between 10 and 48 years of age. Individuals were recorded from a distance of at least 7m while engaged in their natural behaviour. Focal animal sampling was employed [25], involving 15 minutes of continuous video observation of one single animal, with the aim of capturing a clear and complete view of the animal and all its behaviours, including communication. Focal animals were only sampled once a day. Initially focal subjects were chosen on the basis of visibility and ease of pursuit to ensure high-quality recordings. Later in the study period, priority to certain subjects was given in order to homogenize the total focal time across individuals. Thirty-one hours of video data were used in this study.

### Data extraction: the vocal-visual combinations

Subsequent data extraction was carried out on the video/audio recordings using Noldus Observer XT 10 events logging software (http://www.noldus.com/animal-behaviour-research). The annotation of video/audio footage was centered around events of vocal production (N=297). For each of these events, the researcher coded information on both the vocal as well as the visual components of signal production.

Vocalizations were classified according to the call types described in existing chimpanzee repertoires and specific empirical studies [16,26]. Of the ∼13 call types described in the repertoires, this study focused on the seven most commonly produced: grunt, soft hoo, pant bark, pant grunt, pant hoot, scream and whimper. The minimum number of occurrences necessary for a call to be included in the analyses was 5. In the case of the calls “grunt” and “soft hoo”, the existing literature describes different call subtypes, whereby “soft hoo” can be divided into “travel hoo”, “rest hoo” and “alarm hoo”, while “grunt” can refer to “rough grunt” or “general grunt”. Here however, all respective subtypes were lumped into the broad categories of “soft hoo” and “grunt”. Rough grunts and general grunts were collapsed given that our sample only included low-frequency rough grunts, which are acoustically similar to general grunts. High-pitched rough grunts and rare call types did not occur in the available video-audio footage with sufficient frequency to be included in this study. Additional call types that were not observed at least 5 times and therefore not included in the study were the following: bark, waa bark, pant, cough, wraa, laughter, squeak. The number of events observed for the seven call types included ranged from 5 to 98. Chimpanzee vocalizations are often produced in bouts. A bout was defined as a sequence of the same vocalization with pauses shorter than 10s between the individual acoustic elements. A bout was considered terminated when followed by 10s of silence or by the production of a different call type. Bouts constituted single data points. The duration of vocal bouts ranged between 1-62 seconds.

In association with each vocal event, between 1-8 NVBs were recorded. NVBs were only annotated during vocal bouts. A total of 31 different NVB types were recorded in this study. Table 1 provides the full list of NVBs annotated in this study, as well as a description of the behavioural criteria used to assign each NVB type. The NVBs included in this list represent an attempt to illustrate the observable variation in NVB behaviour, and the level of granularity takes into account the risks of an over-representation of NVBs, general feasibility in coding, and complying with inter-observer reliability. Additional measures taken to maximally standardize the annotation procedure can be found in the ESM.

**Table 1.** Full list of NVBs annotated in this study with corresponding behavioural description used to assign NVBs. The term “specific individual” used above refers to the individual who is closest to the signaler.

| NVB name | NVB description |
| --- | --- |
| rest | signaler is lying down or in resting position with chest or back touching the ground |
| sit | signaler sits with bottom touching ground, chest or back are not touching ground |
| get_up | signaler transitions from lying or sitting position to standing or walking |
| stand | signaler is in erect quadrupedal position without movement |
| walk | signaler moves quadrupedally by more than 1 meter |
| run | quadrupedal movement that occurs at a faster pace than normal walking, often gallop-like appearance with both feet in the air at once |
| climb | signaler moves up, down or along the trunk or branch of a tree |
| look_towards | head orientation is shifted toward specific individual by at least 90 degrees resulting in specific individual being in line of sight of signaler |
| look_away | head orientation shifted away from specific individual by at least 90 degrees |
| gaze_upwards | head orientation is shifted towards the canopy/sky |
| gaze_alteration | head orientation changes 3 or more times by approximately 90 degrees |
| turn_body_towards | body orientation changed by at least 90 degrees in direction of specific individual |
| turn_body_away | body orientation is shifted away from specific individual by at least 90 degrees |
| extend_body_towards | signaler moves chest, back or bottom toward a specific individual but legs do not usually move |
| retract_body | signaler's body axis connecting hips to head either changes angle or moves away from specific individual |
| crouch_down | signaler brings bottom, body or shoulders close to the ground |
| present_back | signaler orients back and bottom toward a specific individual by at least 90 degrees |
| arm_reach | arm is fully or partially extended towards a specific individual with or without contact |
| arm_wave | arm performs repetitive back and forth or side to side motion |
| scratch_self | fingers perform loud scratching gesture against any body surface |
| approach | signaler moves in direction of specific individual with 45 degree accuracy on either side |
| embrace | arms or legs are wrapped around a specific individual with degree of surface body contact consisting in at least hand/foot + forelimb |
| chase | signaler runs or climbs quickly after a specific individual in aggressive manner |
| hit | hand or foot is moved aggressively with the intent to make contact with body part of another individual |
| grab_branch | tree branch is grabbed and shaken or dragged along the floor while running or displaying |
| slap_ground | hands or feet are brought violently against the ground to produce a smacking noise, sometimes repeatedly |
| feed | signaler grabs food items and places in mouth, or chews food items already in mouth |
| groom | signaler probes own hair or that of other individual and extracting small particles, using one or both hands |
| play | signaler interacts with another individual via non-aggressive grabbing, biting, chasing, climbing, tickling |
| relaxed_open_mouth_face | open mouth with intermediate separation between upper and lower jaw, while engaged in play |
| scream_face | wide open mouth with maximum separation between upper and lower jaw, lip corners pulled up, teeth bared |

### Data extraction: demographic context of the vocalization

In addition to describing vocal signals and accompanying NVBs, demographic data were annotated for each event. Specifically, identity and sex of the individual were noted and each individual’s age in years was calculated based on the long-term data which includes birth dates for all IDs [24]. Next, dominance ranks were calculated using an Elo-rating method [27,28] based on the long-term data on aggressive interactions and submissive pant grunt vocalizations [29]. Rank scores were calculated every 3 months and ranged between 1-24.

### Inter-observer reliability

To ensure videos were coded reliably, a second independent researcher coded 11% of the events (i.e. 34 events out of 297) and extracted both i) the call type (at least one call for each call type was present in the subset) and ii) non-vocal behaviours (at least one instance of each NVB type was coded in the subset). We calculated a Cohen’s kappa value of 0.82 and 0.88 for vocalisation type and NVB type respectively, indicating excellent levels of agreement in both cases [30].

### Collocation analysis

To generate a vocal-visual signal repertoire based on the communicative events observed, we implemented a collocation analysis in R [31]. This method, originating in the field of linguistics and recently adapted to the study of animal communication, estimates the relative attraction between communicative units, based on how frequently they co-occur in the dataset [14]. In this case, the co-occurrence of a particular vocal signal with a specific visual component was examined. For example, if “grunt” + “arm reach” co-occur, collocation analysis compares the frequency of “grunt + arm reach” with the frequency of all other vocal-visual combinations which contain either “grunt” or “arm reach”. A multiple distinctive collocation analysis tests the association between units via one-tailed exact binomial tests on each possible combination, and the log-transformed results provide an estimate of how exclusively units combine with one another. Ultimately, the test indicates whether each combination happens more or less frequently than expected by chance.

A feature of the communicative events included in this dataset is that one vocal signal commonly co-occurs with more than one NVB simultaneously. For example, a “grunt” vocalization may co-occur with a “sit” posture, a “scratch self” gesture and a “look towards” movement. Our analysis aimed to investigate not only the above-chance occurrence of vocalizations and NVBs individually, but also the association between a given call and multiple NVBs at once. Therefore, a modified collocation analysis was designed to test the association between one call and up to four concomitant NVBs. This threshold of 4 was chosen as 93% of events exhibited between 1-4 NVBs. In order to test associations between vocalizations and NVBs at all levels of combination, each event where >1 NVB occurred was entered into the dataset first with each NVB individually, and then with all possible combinations of two, three and four NVBs given the NVBs present in that event. When such combinations were entered into the data table, this was done while maintaining the two-column structural requirement of collocation analyses as shown in Table 2.

**Table 2.**
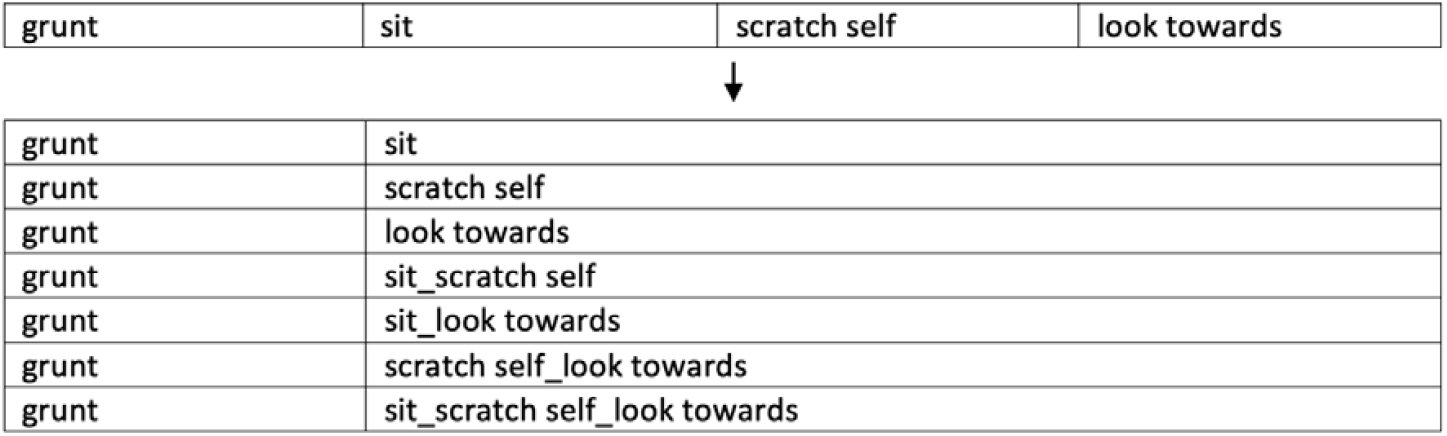
Illustration of procedure for entering each communication event into a suitable dataset for implementing the multiple-NVBs collocation analysis.

|  |  |  |  |
| --- | --- | --- | --- |
| grunt | sit | scratch self | look towards |

|  |  |
| --- | --- |
| grunt | sit |
| grunt | scratch self |
| grunt | look towards |
| grunt | sit_scratch self |
| grunt | sit_look towards |
| grunt | scratch self_look towards |
| grunt | sit_scratch self_look towards |

### Statistical analyses: demographic and call-related drivers of NVB production

To examine variation in the number of NVBs produced alongside vocalizations as a function of demographic variation and call characteristics (i.e. call type and call duration), we performed a generalized linear mixed model (GLMM) with a negative binomial error structure and log link function using the glmmTMB function, glmmTMB package in R. We modeled the number of NVBs produced per event as a numerical integer response variable. As demographic predictors, we fitted age (years) as a second-order polynomial, sex as a binary categorical variable (M/F) and rank as a numerical integer. As call-related predictors, we fitted call type as a 7-level categorical variable, and duration of call bout (seconds) as a numerical predictor. Given that the effect of call type and duration may not be independent, an interaction term was fitted between these predictors. Individual identity was fitted as a random factor to account for multiple events from single individuals.

We first compared the full model including all predictors and random effects with a null model which was identical in structure minus the predictors, for which we report a likelihood ratio test (chi-squared statistic and p-value). We ascertained the relative contribution of each variable to the model by comparing the full model to a reduced model lacking each individual predictor in turn. We then report chi-squared values of likelihood ratio tests regarding the effect of each individual predictor, as well as p-values using a 95% significance threshold.

Model assumptions were checked using the DHARMa package in R. The model was not found to exhibit overdispersion (nonparametric dispersion test P = 0.74), no outliers were detected (P = 0.4) and visual inspection of the Q-Q plots confirmed normality (Kolmogorov-Smirnov test: P = 0.77).

## Results

### Vocal-visual repertoire via collocation analysis

Following collocation analyses, 108 combinations of one vocal signal and between 1-4 NVBs were found to co-occur significantly more frequently than expected by chance (all p values <0.05). The number of significant combinations varied between call types: for example, four combinations were documented for the “pant bark” call, six for the “scream”, 11 for the “whimper”, 16 for the “soft hoo”, 22 for the “pant grunt”, 24 for the “pant hoot” and 25 combinations for the “grunt” call. Of the 31 NVB types present in the raw data, 21 featured in significant combinations with vocal signals. Eighteen out of these 21 NVB types (i.e. 86%) were recombined productively across multiple call types. The full set of significant combinations which constitute the vocal-visual repertoire is presented in Tables 3 and 4.

**Table 3.**
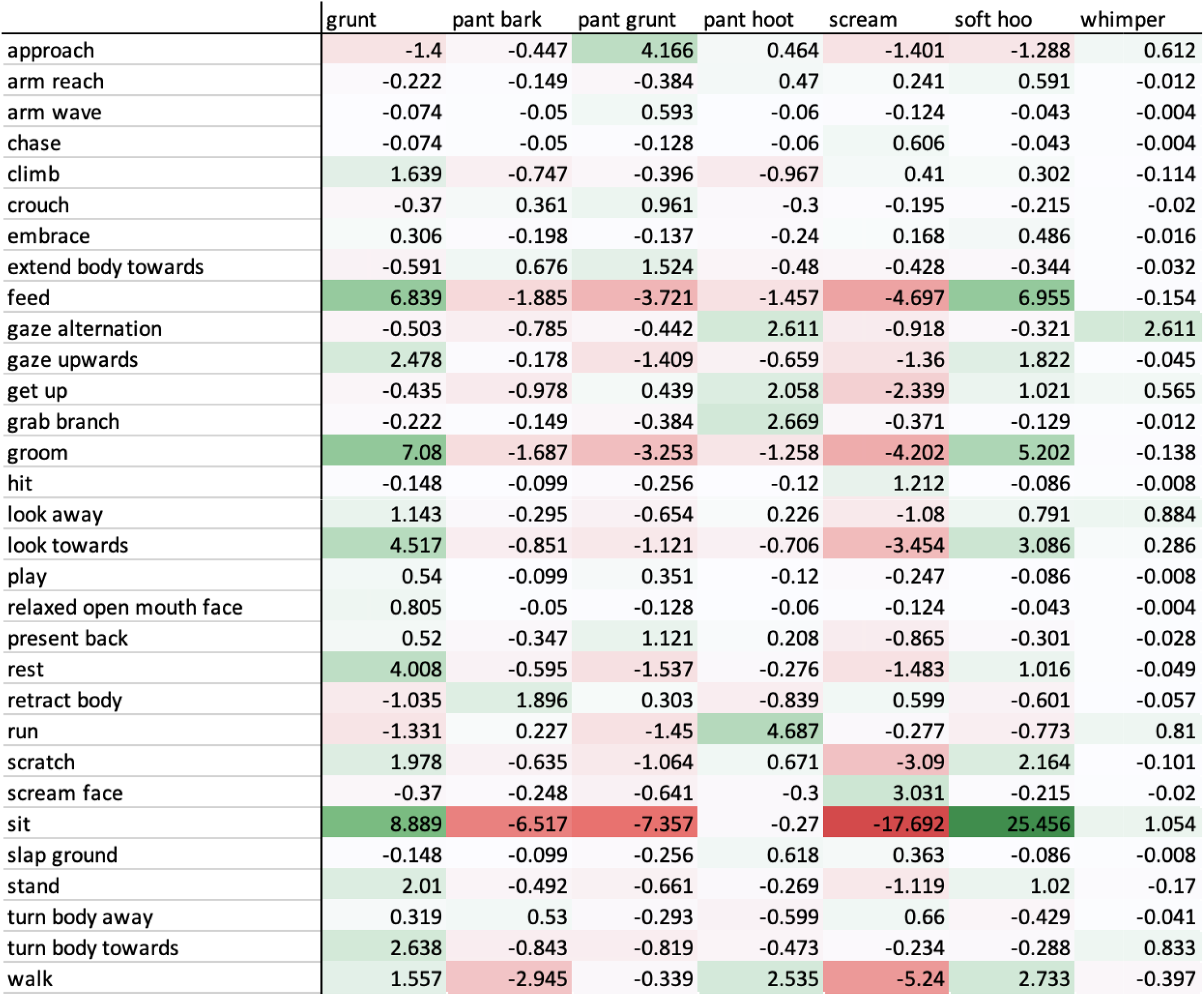
List of 31 single NVBs and 7 call types included in this analysis. Colour codes denote strength of attraction/repulsion between NVBs and each call type: darkest green = strongest attraction, darkest red = strongest repulsion. All values above 1.3 represent co-occurrence at above-chance level with 95% confidence interval, while values below -1.3 represent significant repulsion between collocates.

**Table 4.** All combinations of call type and NVBs that were found to co-occur significantly more frequently than expected by chance.

| grunt | pant hoot | pant grunt | soft hoo | whimper | scream | pant bark |
| --- | --- | --- | --- | --- | --- | --- |
| climb | approach_get up_sit | approach | feed | approach_get up_look tow_run | look towards_scream face | extend body_look to |
| climb_feed | approach_get up_sit_walk | approach_extend body | feed_sit | approach_get up_run | scream face | look tow_retract bo |
| feed | gaze alternation | approach_extend body_getup | feed_sit_stand | approach_look tow_run | scream face_stand | look tow_retract bo |
| feed_sit | gaze alternation_get up | approach_extend body_get up_walk | feed_stand | approach_run | scream face_turn body tow | retract body |
| feed_walk | gaze alt_get up_sit | approach_extend body_walk | gaze upwards | gaze alternation | stand_turn body towards |  |
| gaze upwards | gaze alt_get up_sit_walk | approach_get up | gaze upwards_sit | gaze alternation_sit | stand_turn body tow_walk |  |
| gaze upwards_sit | gaze alt_get up_walk | approach_get up_walk | groom | gaze alt_sit_turn body towards |  |  |
| groom | gaze alternation_scratch | approach_look towards_walk | groom_sit | gaze alternation_turn body tow |  |  |
| groom_sit | gaze alt_scratch_sit | approach_present back | look towards | get up_look towards_run |  |  |
| look away_stand_walk | gaze alternation_sit | approach_present back_walk | look tow_scratch self | get up_run |  |  |
| look towards | gaze alternation_sit_walk | approach_walk | look tow_scratch_sit | look towards_run |  |  |
| look towards_rest | get up | climb_gaze alternation_stand | look towards_sit |  |  |  |
| look towards_sit | get up_sit | climb_gaze alternation_stand_walk | scratch self |  |  |  |
| look tow_sit_turn body towards | get up_sit_walk | climb_gaze alternation_walk | scratch self_sit |  |  |  |
| look tow_sit_turn body tow_walk | get up_walk | climb_stand | sit |  |  |  |
| look towards_turn body towards | grab branch | climb_stand_walk | walk |  |  |  |
| rest | grab branch_run | climb_walk |  |  |  |  |
| scratch self | grab branch_run_walk | extend body |  |  |  |  |
| scratch self_sit | grab branch_walk | extend body_get up |  |  |  |  |
| sit | run | extend body_get up_walk |  |  |  |  |
| sit_turn body towards | run_walk | extend body_walk |  |  |  |  |
| sit_turn body towards_walk | scratch self_walk | present back_walk |  |  |  |  |
| stand | sit_walk |  |  |  |  |  |
| turn body towards | walk |  |  |  |  |  |
| walk |  |  |  |  |  |  |

### Demographic and call-related drivers of NVB production

Our GLMM analysis indicated that the full model, including all predictors, explained significantly more variation in the response variable compared to a null model (χ^2^_16_ = 38.96, p = 0.001). Likelihood ratio tests revealed that there was no significant main effect of age (χ^2^_2_ = 1.39, p = 0.49), sex (χ^2^_1_ = 1.25, p = 0.26) or rank (χ^2^_1_ = 1.29, p = 0.25) on the number of NVBs produced per vocalization. However, there was a significant interaction between call type and duration (χ^2^_6_ = 19.68, p = 0.003), such that the effect of duration on the number of NVBs differed between call types. Longer call duration was associated with more NVBs in “pant grunt”, “pant hoot” and “soft hoo” calls, while no such effect was observed in the other call types. Overall, the “pant grunt” call was produced in association with the most NVBs while the “scream” was associated with the fewest, as shown in Figure 1.

**Figure 1.**
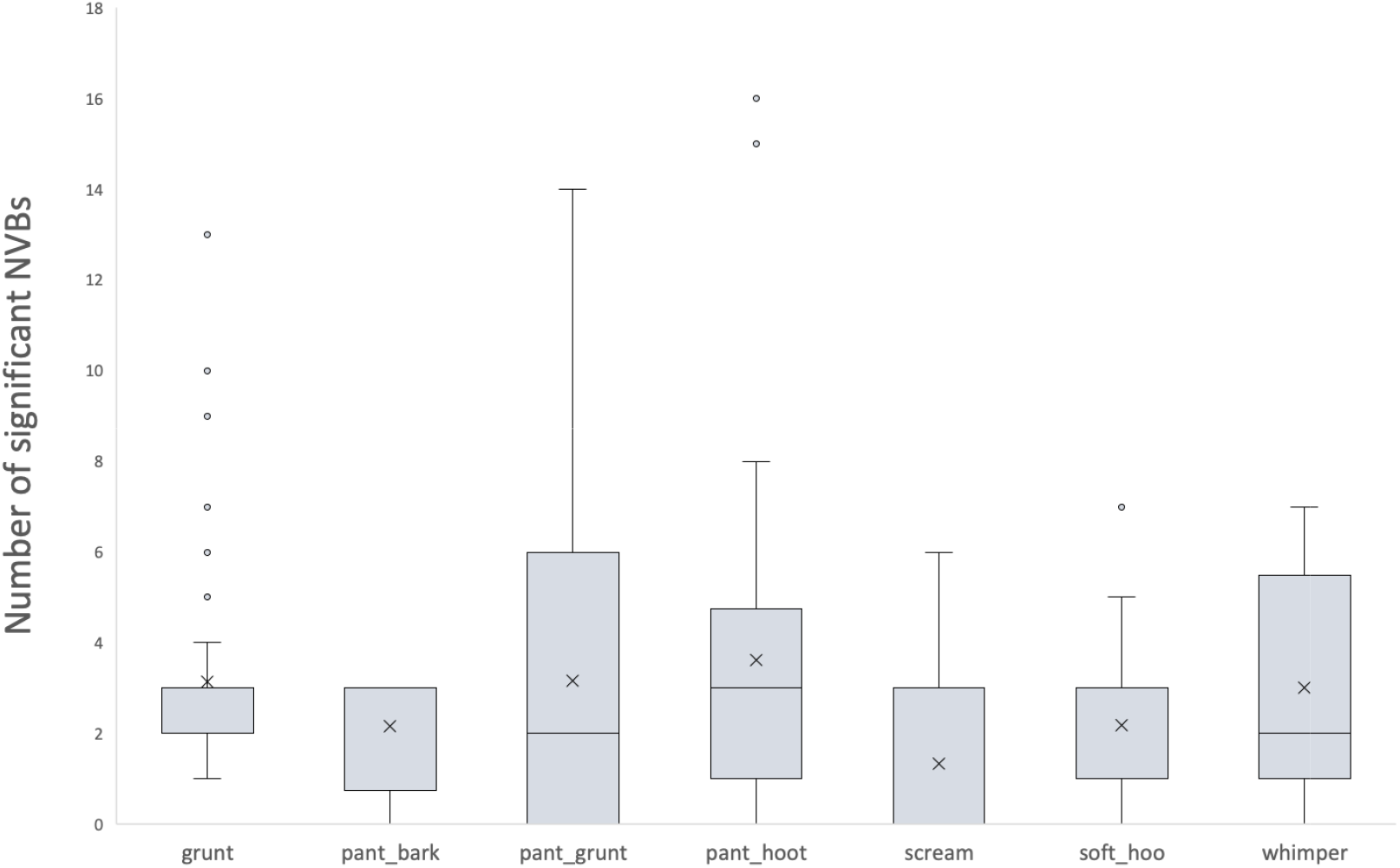
Raw data illustrating variation in the number of significant NVBs produced in association with the different call types analysed in this study. Crosses represent means for each call type.

## Discussion

By systematically observing naturally occurring communication events, we show that chimpanzees combine their vocal signals with a wide range of body movements, postures, gestures and facial expressions, collectively referred to here as non-vocal behaviours (NVBs). More than 100 such combinations of vocal and visual components occur more frequently than expected by chance, indicating a strikingly diverse repertoire of vocal-visual combinations. Some NVBs are used productively across multiple call types, yet each call type is associated with its own set of single and combined NVBs. When a vocalization is produced, the number of accompanying NVBs increases with call duration, but this effect is conditional on call type, such that longer vocalization events are associated with a greater number of NVBs in some call types but not in others. However, the number of NVBs associated with vocal production is not influenced by age, sex, or rank.

Given the findings of the collocation analysis, it appears that sub-adult and adult chimpanzees have access to a highly diversified repertoire of combined visual and vocal components. Although the constrained vocal repertoire of chimpanzees [19,20] might suggest a limited capacity for information transfer, the productive use of accompanying NVBs instead reveals a high potential for refining the meaning of the limited range of available calls. Indeed, the ∼100-strong repertoire of combinations reported in this study highlights the potential for extensive and nuanced information transfer between communicating chimpanzees. A fundamental implication of this investigation is that unimodal approaches to primate communication, which analyze vocal or visual components separately, result in a drastically oversimplified picture of flexibility in signal production. A multi-modal approach is therefore crucial to accurately representing the communicative abilities of non-human primates [5,6], as well as for offering a faithful illustration of real-life communicative exchanges.

Chimpanzee social life is characterized by a wide variety of interactions, each of which is typically mediated by communication. Thus, it is likely that the diverse repertoire of combined vocal and visual components identified here plays a key role in supporting the demands of a chimpanzee’s daily social life [32,33]. It is unknown whether chimpanzee signalers voluntarily combine vocal signals with all of the NVBs reported in this study, nonetheless, chimpanzee receivers may rely on the integration of all the vocal and visual components in order to guide their own adaptive behavioural response [34]. Confirming this hypothesis requires further investigation into how NVBs are perceived by receivers and their potential role in the disambiguation of meaning. Recent developments which combine insights from linguistics and animal behaviour offer valuable theoretical frameworks and empirical toolkits for addressing the meaning of signal components empirically in nonhumans [35]. One fruitful method involves a systematic analysis of behavioural reactions to signals as a function of signal type [36]. This method could be applied to the wide range of vocalization and NVB combinations highlighted in this study, offering critical insights into the meaning of chimpanzee vocal-visual combinations. A further promising avenue of investigation is to infer which cues are most salient to recipients for meaning disambiguation, using measures of attentional bias. The application of eye-tracking technology in captive great apes, for example, has enjoyed a recent surge of advances, bringing this goal confidently within reach [37].

Our study also investigated the variation in the number of NVBs produced per vocalization as a function of individual demographic attributes such as age, sex and rank. However, males and females did not differ in the number of NVBs produced, nor was the observed variation explained by age or rank. A possible implication of this result is that combinatoriality across modalities may serve a very general function such as that of meaning refinement, which is critical irrespective of demographic status. Replicating this work in other communities of chimpanzees would prove useful for establishing the universality of this finding. Indeed, it remains possible that a population which experiences different ecological or social pressures, may display more pronounced demographic patterns in NVB production than those observed here.

In conclusion, our findings reveal a hitherto unappreciated diversity of vocal-visual combinations in the communication system of wild chimpanzees, though follow-up behavioural observations and experimental work are key to unpacking the function and meaning of such combinations. Nonetheless, the extent and variety of non-random vocal-visual combinations described here broadens our appreciation of the potential combinatorial information available to receivers in our closest-living relative. Furthermore, ∼90% of the visual components of communicative exchanges observed in this study were shown to be produced in association with multiple call types. In line with previous work, this is suggestive that multi-modal signals represent combinatorial structures, of which vocal and visual components constitute the building-blocks, as opposed to holistic units [38]. By virtue of our phylogenetic proximity to chimpanzees, the range of vocal-visual combinations presented here also informs our understanding of the communicative behaviour of our hominin ancestors, suggesting a capacity for complex multi-modal signaling that predates the language faculty and may have played a role in scaffolding language evolution [39-42].

## Supporting information

Electronic Supplementary Material

## Acknowledgements

We are grateful to the directors of Kibale Chimpanzee Project for permitting and supporting us to carry out this research on the Kanyawara community of chimpanzees. We are also thankful to the KCP field manager Emily Otali and the KCP field assistants, Dan Akaruhanga, Seezi Atwijuze, Sunday John, Richard Karamagi, James Kyomuhendo, Francis Mugurusi, Solomon Musana and Wilberforce Tweheyo, for their valuable assistance and support in the field. We thank Piera Filippi for her constructive comments. This project was funded by a Leakey Foundation General Grant to C.W. and the NCCR Evolving Language. We appreciate the permission of the Uganda National Council for Science and Technology, the President’s Office and the Uganda Wildlife Authority for us to carry out this study in Uganda.

## Data and materials availability

All data are available in the supplementary materials.

