## Supplementary material for "Vocal-visual combinations in wild chimpanzees": Electronic Supplementary Material

### Electronic Supplementary Material for "Vocal-visual combinations in wild chimpanzees" by Mine et al. 2023

**This PDF file includes:**

Table S1: additional notes for annotation

**Other Supplementary Materials for this manuscript include the following:**

Data S1: full datasets used in this study

**Additional notes used for standardization of annotation**

| applies to | description of procedure |
|---|---|
| general | If a movement or cue begins before the vocalization, but the vocalization is initiated before the movement is finished, then the movement is still annotated |
| general | NVBs produced during the silent pauses of less than 10s within a vocal bout are still annotated |
| looking behaviour | If the signaler changes gaze orientation by 90 degrees and then resumes original gaze position, this is annotated as both look away + look towards |
| looking behaviour | When 3 or more gaze shifts of 90 degrees are performed, all other gaze shifts are removed and substituted by "gaze alternation" |
| body movement | "Turn body away" does not imply "look away" because "look away" is a 90 degree turn of the head relative to the orientation of the body |
| body movement | The NVB "approach" implies orientation of the body towards another individual, thus the NVB "turn body towards" is not annotated in addition to "approach" |
| body movement | When the body is turned away from the receiver by 90 degrees as a consequence of movement (e.g. walk, run, climb), this is still counted as "turn body away" |
| aggression | When "chase" is coded, "approach" is not coded, to avoid over-representation of cues that effectively refer to the same behaviour |
| aggression | If there is an attempt to hit but without contact, this is still coded as "hit" |

**Table S1.** In addition to the coding scheme presented in the main text of the manuscript, we report here additional measures employed during the annotation of vocal and visual components of communicative interactions. These measures were adopted with the aim of rendering the annotation procedure as standardized as possible, in order to maximize inter-observer reliability.

**Data S1. (separate file)**
Vocal-visual combination datasets including all data used for:

- Annotation of vocal and visual components of communicative interactions
- Collocation analysis
- GLMM on demographic variation in NVB production

These data can be found at the following link:
https://osf.io/gc854/
